## Supplementary Figures for "A thousand-genome panel retraces the global spread and climatic adaptation of a major crop pathogen"

Alice Feurtey et al.

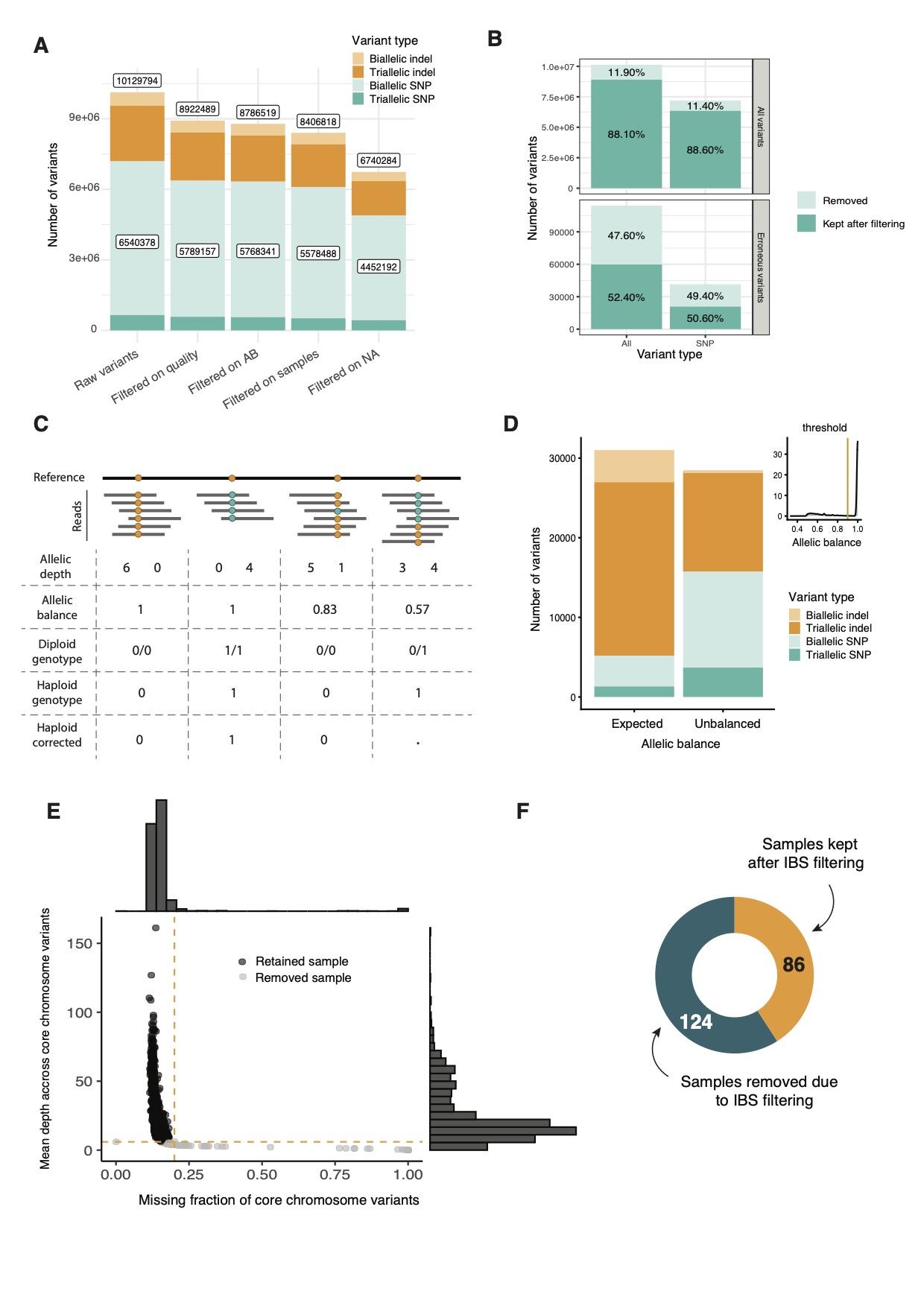
**Figure S1: Short variants filtering procedure.**

A. Number of short variants retained after each filtering step from the raw variant calling of GATK to the final filtered dataset.
B. Number and proportions of variants filtered by hard filtering on quality and depth per position. Top panel: all variants. Bottom panel: only erronous variants detected through the analysis of replicate resequencing sets of the same isolate.
C. Schematic representation of the allelic balance (AB) and the associated filtering. The schematic shows a threshold of 0.8 as an example (we used 0.9 in our analyses). Four different scenarios of allelic depth, allelic balance, called diploid and haploid genotypes both pre- and post-filtering. The dot represents a genotype set to missing data during variant calling.
D. Number of variants of each type (colors as in panel A) for both variants found with a typical AB and with unbalanced AB, based on the threshold illustrated in the density plot of the upper right corner of the panel.
E. Filtering at the sample level based on mean depth of coverage and missing data proportions on core chromosomes.
F. Number of samples removed and kept amongst the 210 isolates with high relatedness to at least one other isolate.

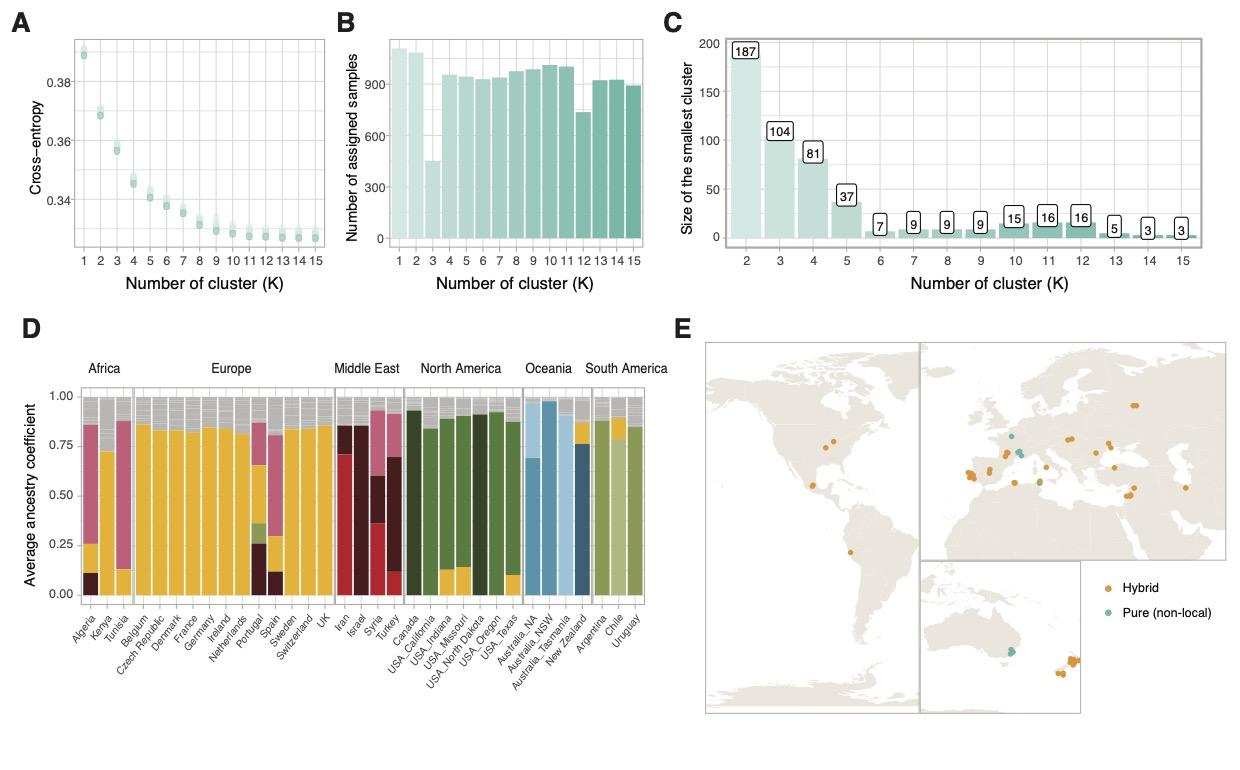

**Figure S2: Genetic clustering of 1109 *Zymoseptoria tritici* isolates and hybrid detection.**

A. Cross-entropy for each number of clusters (K) as assessed by the R package *LEA*.
B. Number of isolates uniquely assigned to one cluster per K.
C. Number of isolates assigned to the smallest cluster.
D. Barplots for the best K (as in panels A-C) per country and state (USA). Color represent the main clusters and correspond to the clusters shown in Figure 1.
E. Map representing the isolates identified either as hybrid or as fully belonging to a cluster that is not the typical local cluster for the country.

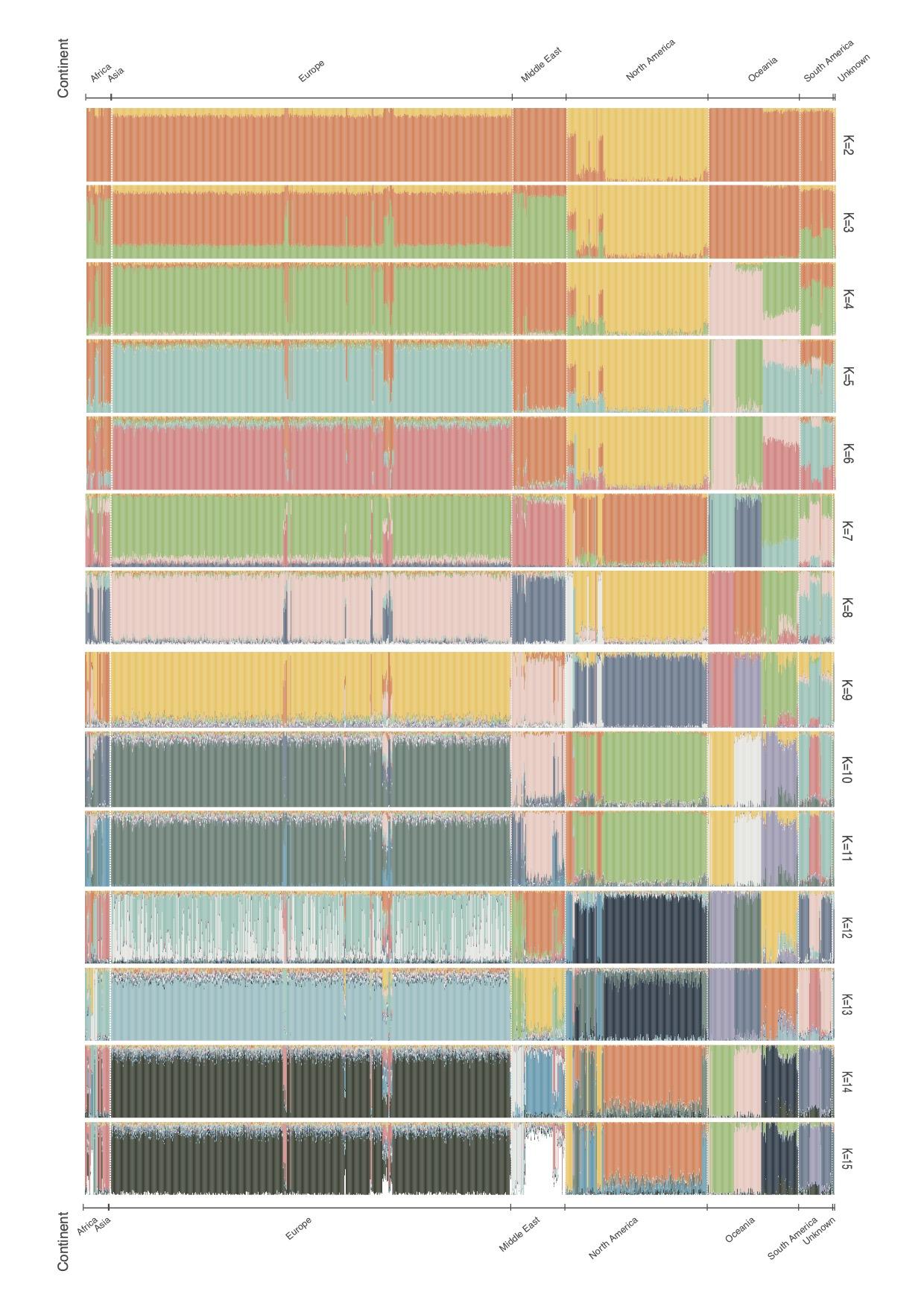

**Figure S3: Genetic clustering of 1109 *Zymoseptoria tritici* isolates showing per-isolate barplots with cluster assignments.** Barplots for the best replicate for each K (as determined by cross-entropy). Barplots are shown for each analyzed value of K. Each vertical bar represents an isolate and the clusters are represented by different colors. Isolates are sorted by continent and per country.

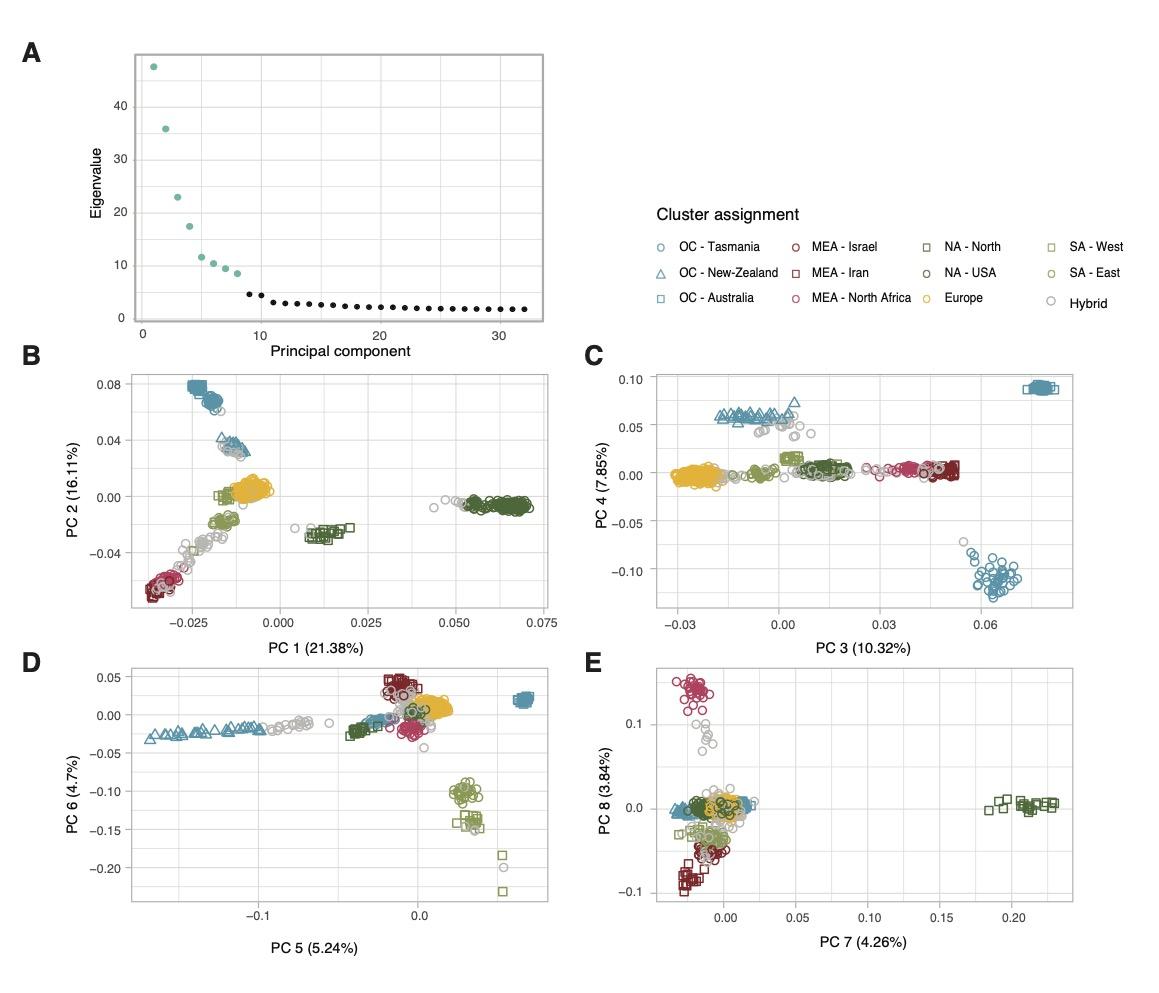

**Figure S4: Principal component analysis based on a subset of the short variant dataset.**

A. Eigenvalues per principal component (PC). The PCs visualized in panels B-E are shown in blue.

B - E. Pairs of PC1-8 with colors and shapes identifying the 11 clusters as defined by the clustering analysis. Genotypes shown in grey could not be attributed fully to any cluster and are thus classified as inter-cluster hybrids.

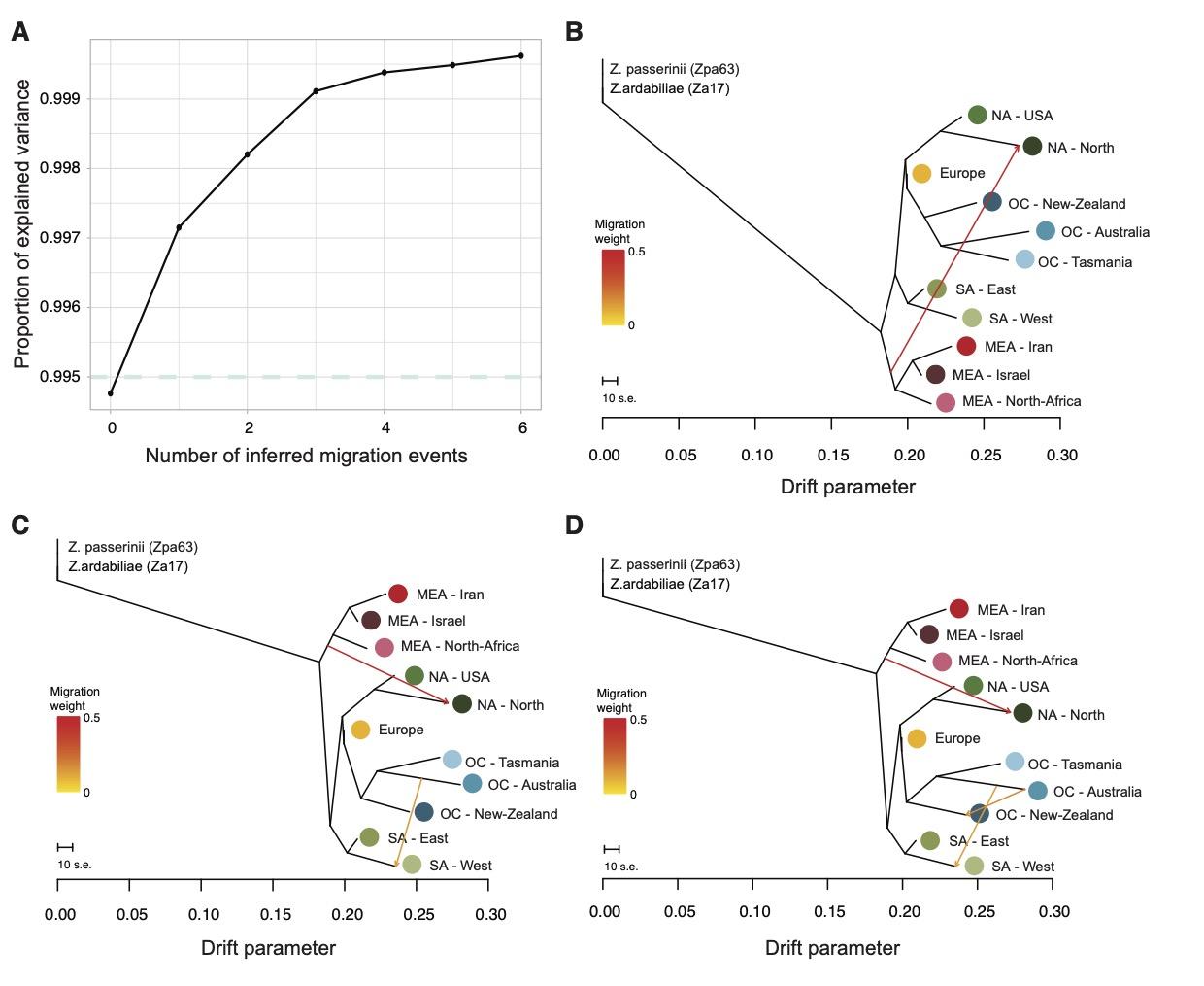

**Figure S5: Population trees (Treemix analysis) with various numbers of inferred migration events**

A. Proportion of explained variance assuming between 0-6 inferred migration events.
B. Tree representation for 1 migration event.
C. Tree representation for 2 migration events.
D. Tree representation for 3 migration events.

Figure S6: Isolation-by-distance per continent assessed by relatedness and geographic distance between sampling locations.

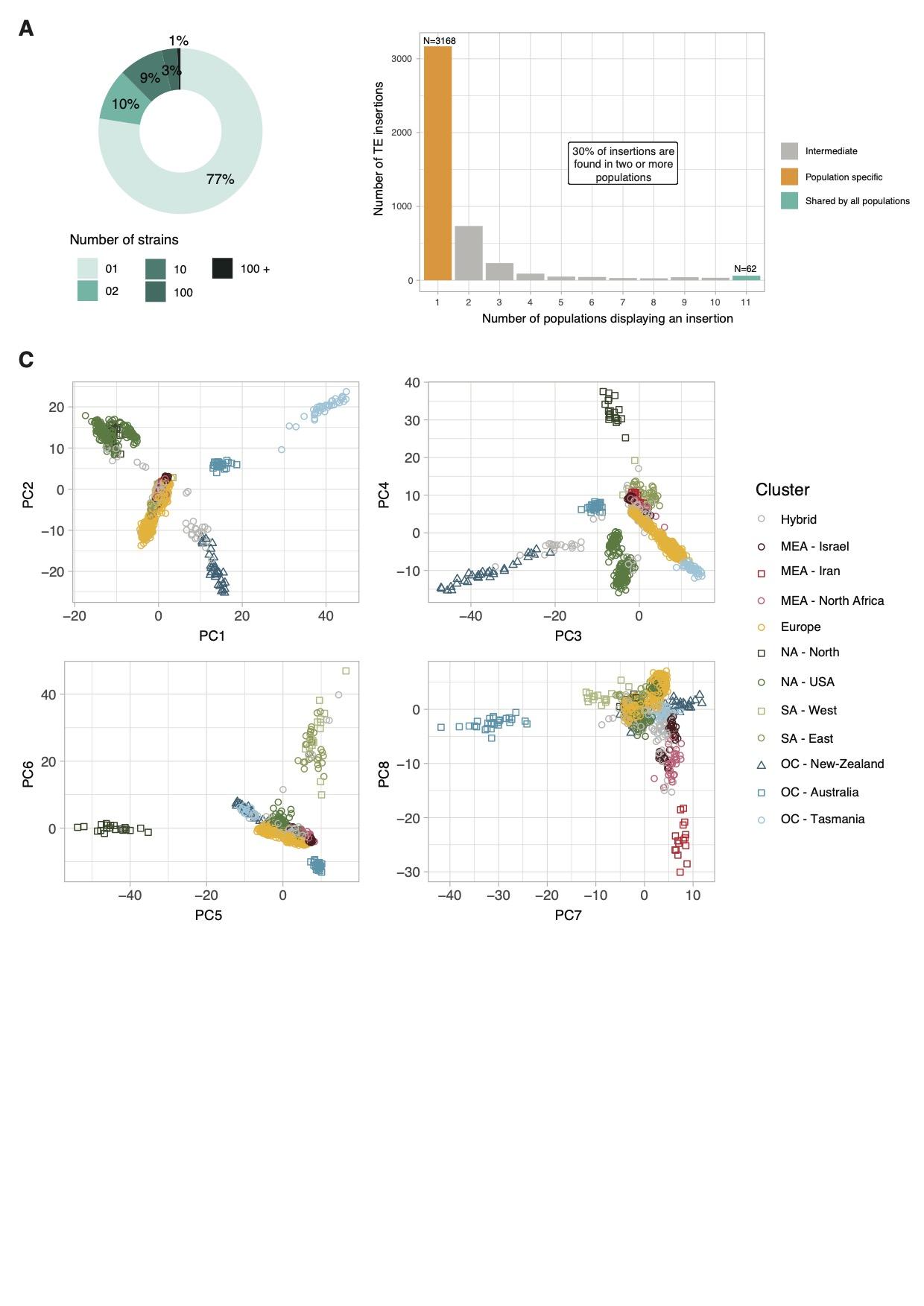

**Figure S7: Transposable element insertion polymorphisms (TIPs).**

A. Proportion of TIPs shared by different numbers of isolates.
B. Number of TIPs found uniquely in one population (orange), found in a subset of populations (grey) or all populations (green).
C. Principal component analysis (PC 1 to 8) based on TIPs shared by at least 10 isolates. The shape and colors of the dots represent the genetic clusters identified based on the short variant polymorphisms.

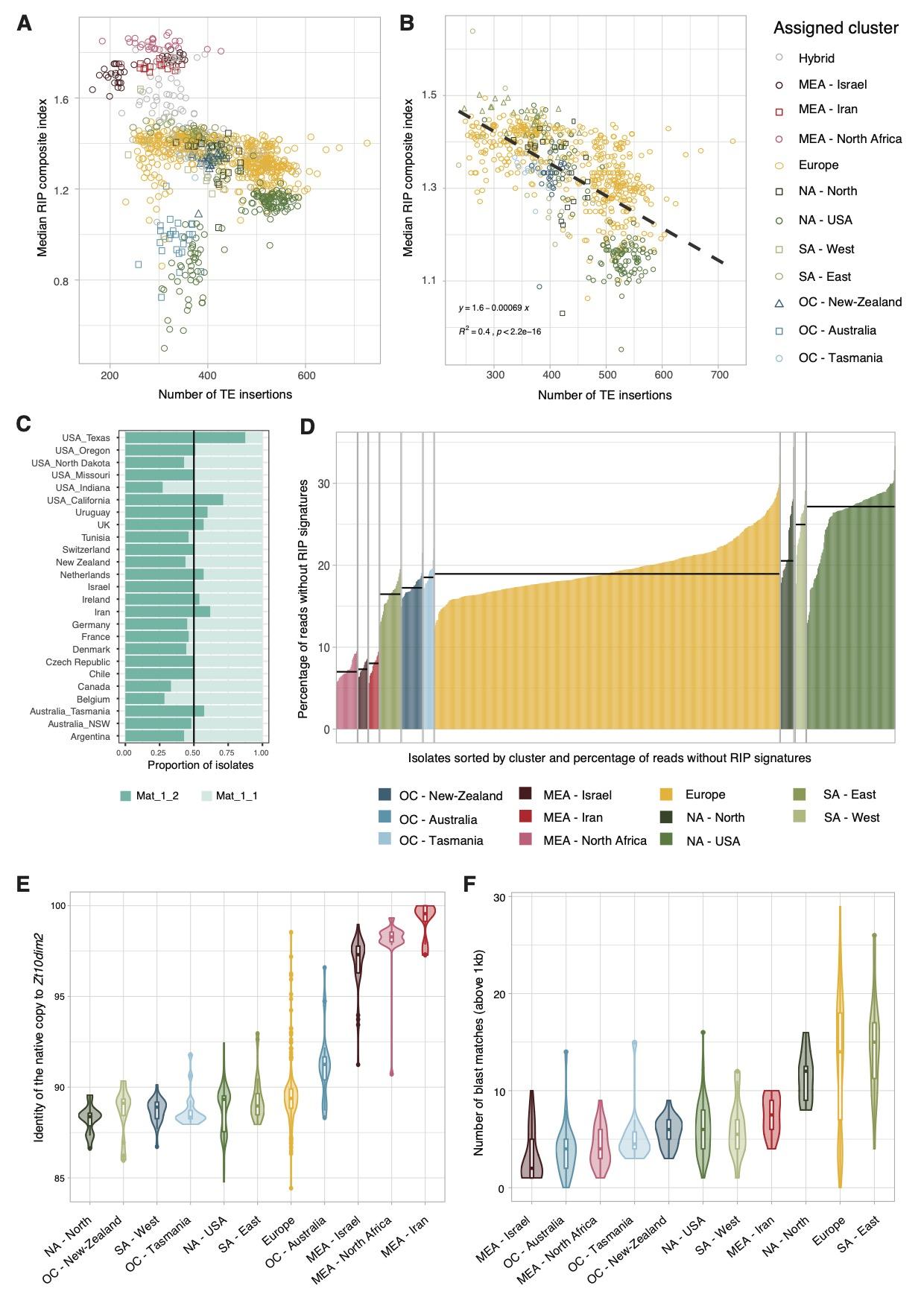

**Figure S8: Patterns of repeat-induced point mutations (RIP) across populations**

A. Mean RIP composite index in TE-mapped reads and TE insertion counts in the global collection of genomes. Colors and shape of dots in panels A and B correspond to the genetic clusters identified based on short variant polymorphisms.
B. Correlation between the RIP composite index in TE-mapped reads and TE insertion counts (excluding the Middle-East/Africa clusters).
C. Proportions of isolates of different mating types (carrying either Mat_1_2 and Mat_1_1) per country/state. The black line represents balanced proportions among mating types (*i.e.* 0.5).
D. Percentage of TE-mapped reads per isolate showing no evidence of RIP (based on a RIP composite index threshold of 0.5). The median percentage per genetic cluster is represented as a horizontal black line.
E. Violin plots and boxplots representing the identity of the native copy of dim2 per cluster (based on a blastn search, and using the flanking genes to identify the native locus).
F. Violin plots and boxplots representing the identity (%) and number of long BLASTn matches (>1 kb) of the functional copy of *dim2* in draft genome assemblies of each isolate grouped by genetic cluster.

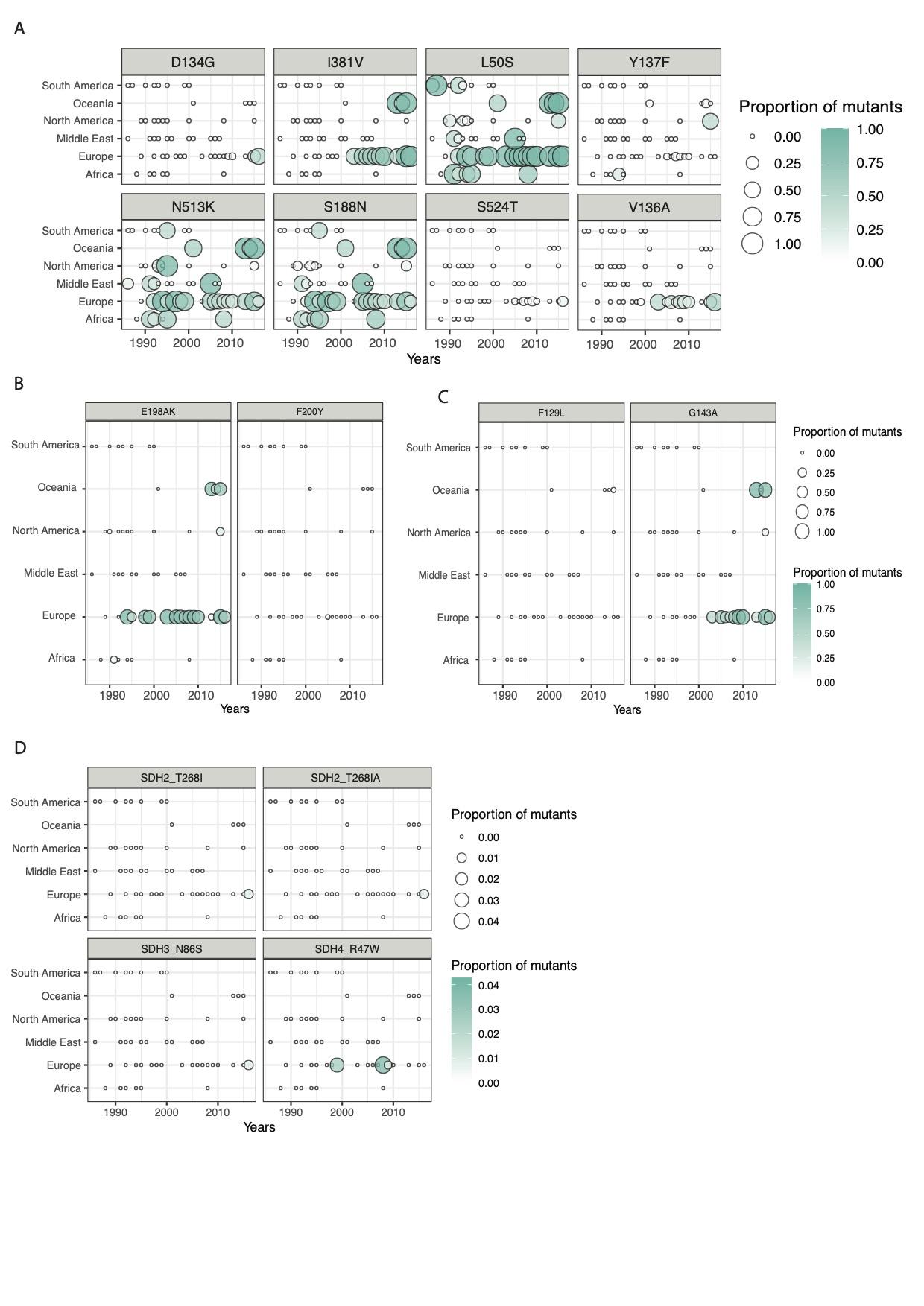

**Figure S9: Survey of mutations known to be associated with resistance to fungicides or observed in natural populations in gene known to be involved in resistance**

A. Mutations in the cyp51 gene, known to be involved in azole resistance.
B. Mutations in the beta tubulin gene, known to be involved in resistance to benzimidazole fungicides.
C. Mutations in the mitochondrial gene cytb, known to cause resistance to Quinone outside inhibitors fungicides.
D. Mutations in the SDH genes, known to be involved in resistance to SDHI fungicides.

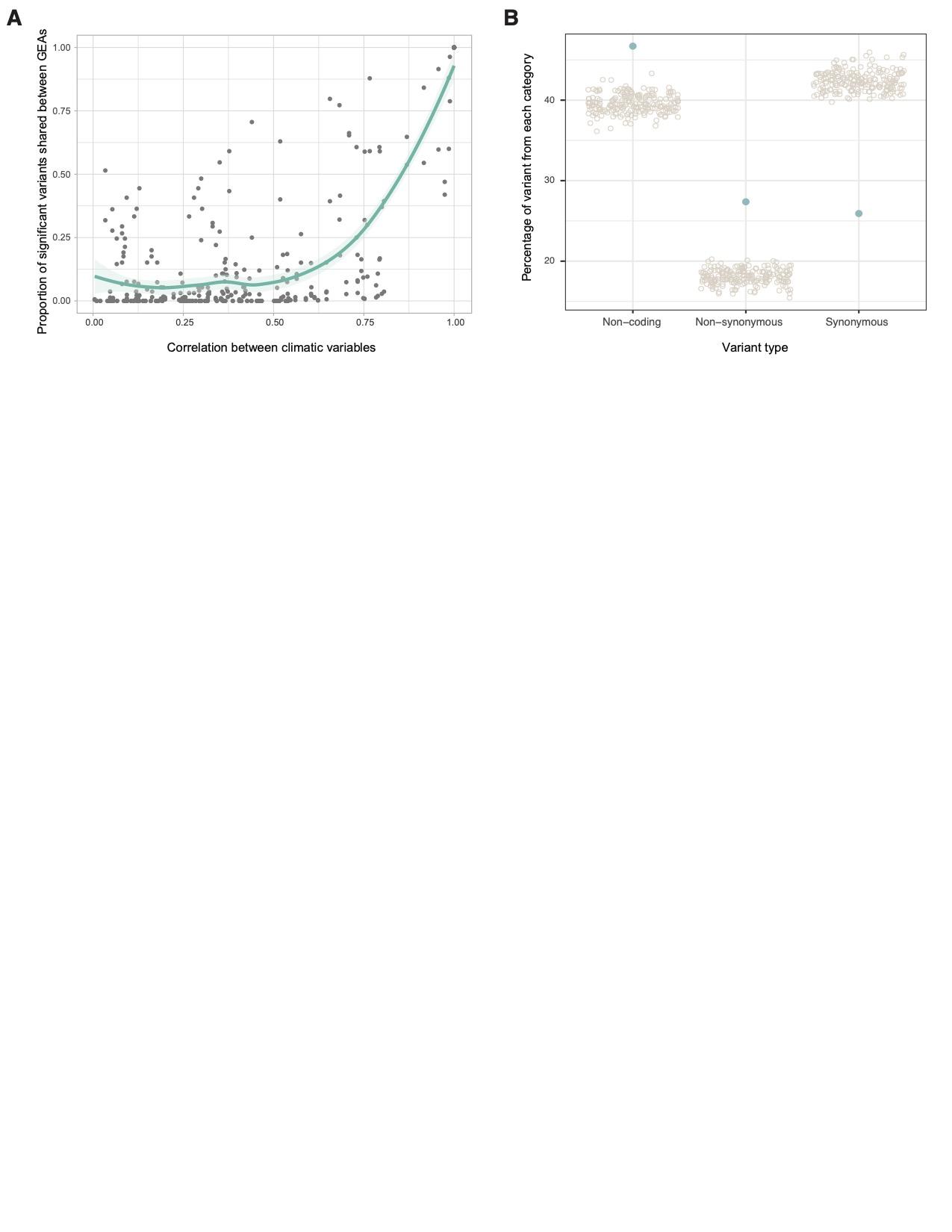

**Figure S10: Genome-environment association mapping: similarity between environmental variables and impact of the significantly associated variants on the predicted protein sequences.**

A. Dot plot showing the correlation between bioclimatic variables and the proportion of variants detected as significantly associated to both of the bioclimatic variables. The green curve was obtained with the loess method, which fits a polynomial using local fitting, span = .75.
B. Comparison between the percentage of variants in non-coding regions, or having synonymous or non-synonymous effects (in green) compared against 200 randomly sampled short variants (with a MAF > 0.01 to match the threshold used for the GEA).

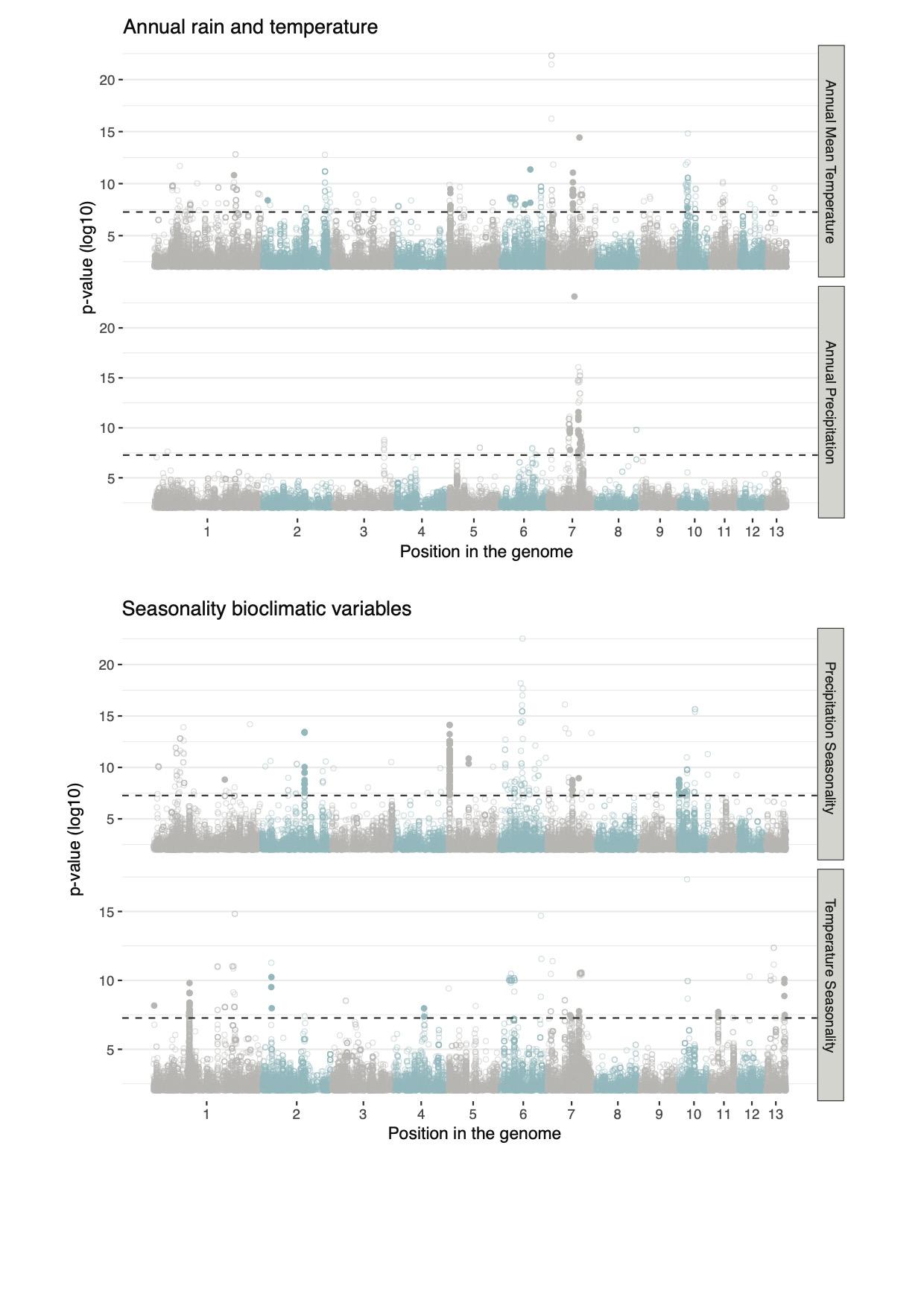

**Figures S11: Manhattan plots of the GEA for bioclimatic variables related to annual rain and temperature levels and to seasonality.**

The x-axis represents the genomic position, each dot represents a short variant with the colors showing alternating chromosomes. The black dashed line is the Bonferonni threshold. Each variant above the threshold is either a filled circle to show a minor allele frequency (MAF) above 5% or an empty circle for lower MAF.

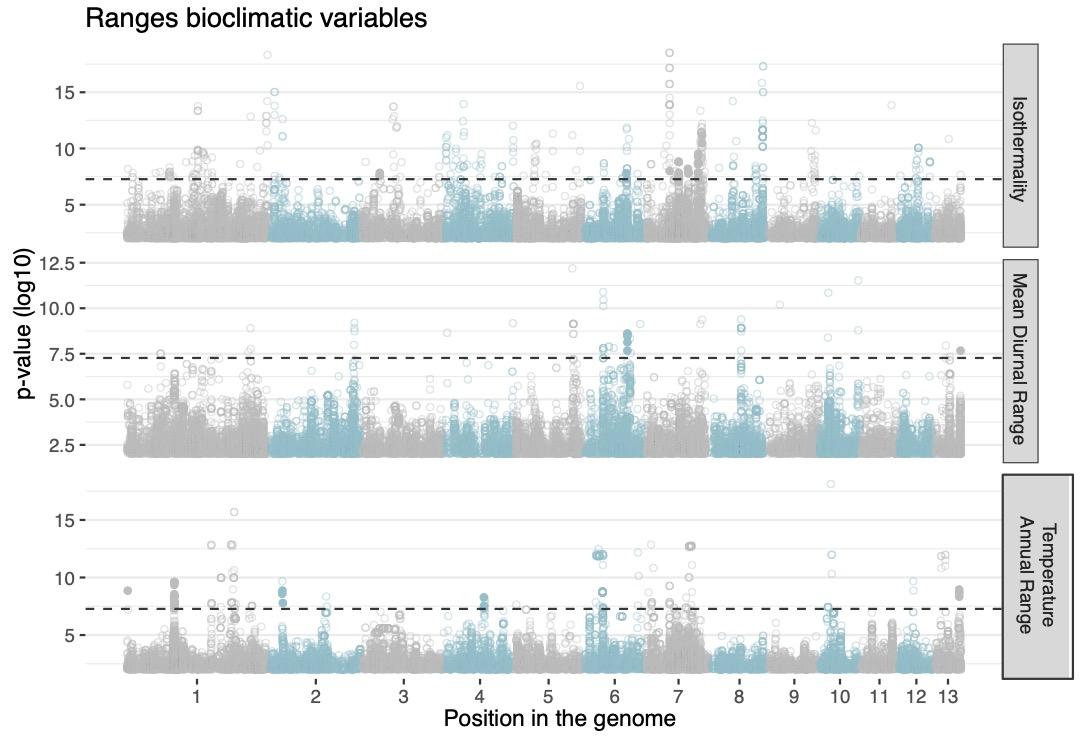
**Figures S12: Manhattan plots of the GEA for the isothermality, the mean diurnal range and the annual temperature range.**

The x-axis represents the genomic position, each dot represents a short variant with the colors showing alternating chromosomes. The black dashed line is the Bonferonni threshold. Each variant above the threshold is either a filled circle to show a minor allele frequency (MAF) above 5% or an empty circle for lower MAF.

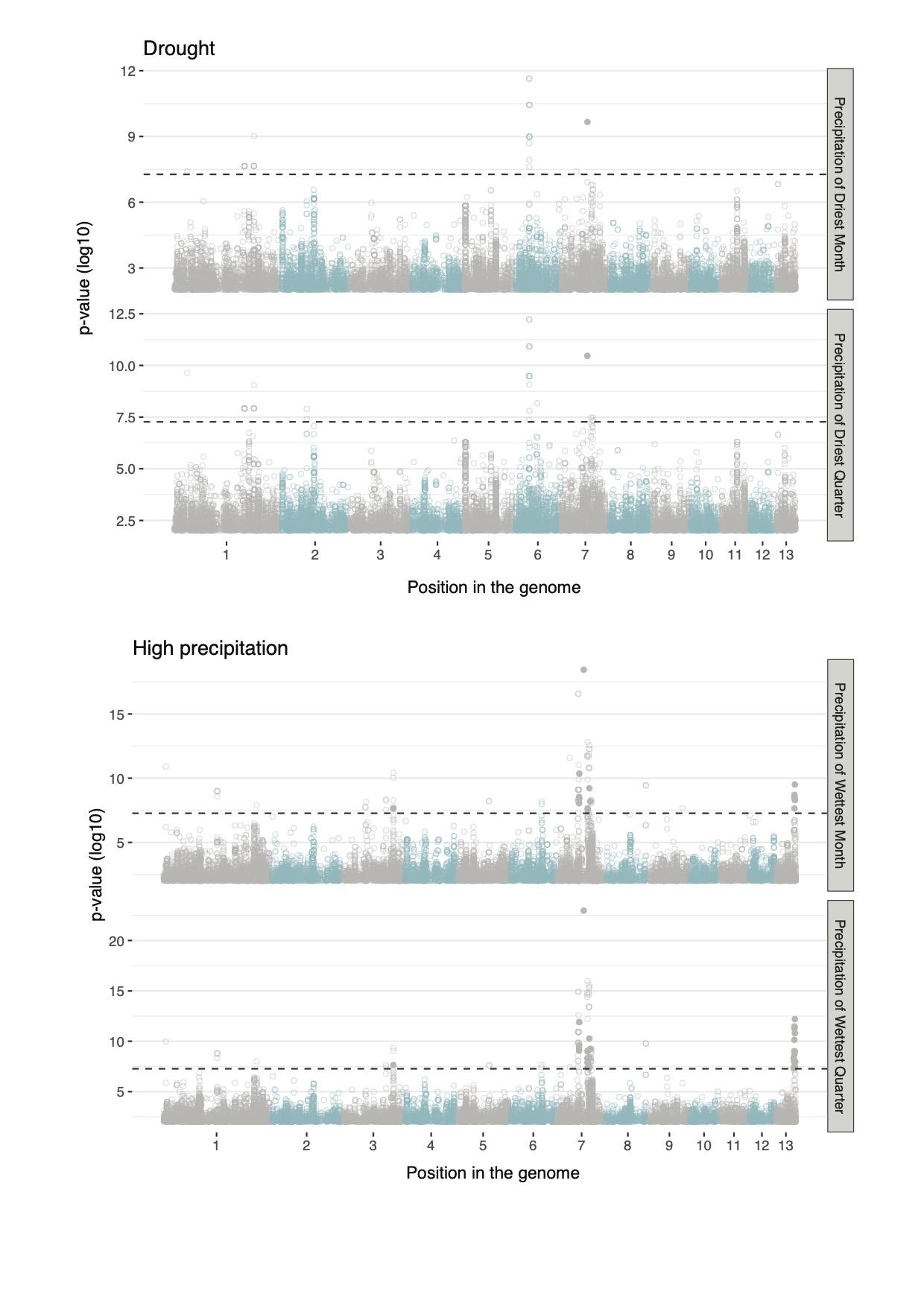

**Figures S13: Manhattan plots of the GEA for high and low precipitation variables.**

The x-axis represents the genomic position, each dot represents a short variant with the colors showing alternating chromosomes. The black dashed line is the Bonferonni threshold. Each variant above the threshold is either a filled circle to show a minor allele frequency (MAF) above 5% or an empty circle for lower MAF.

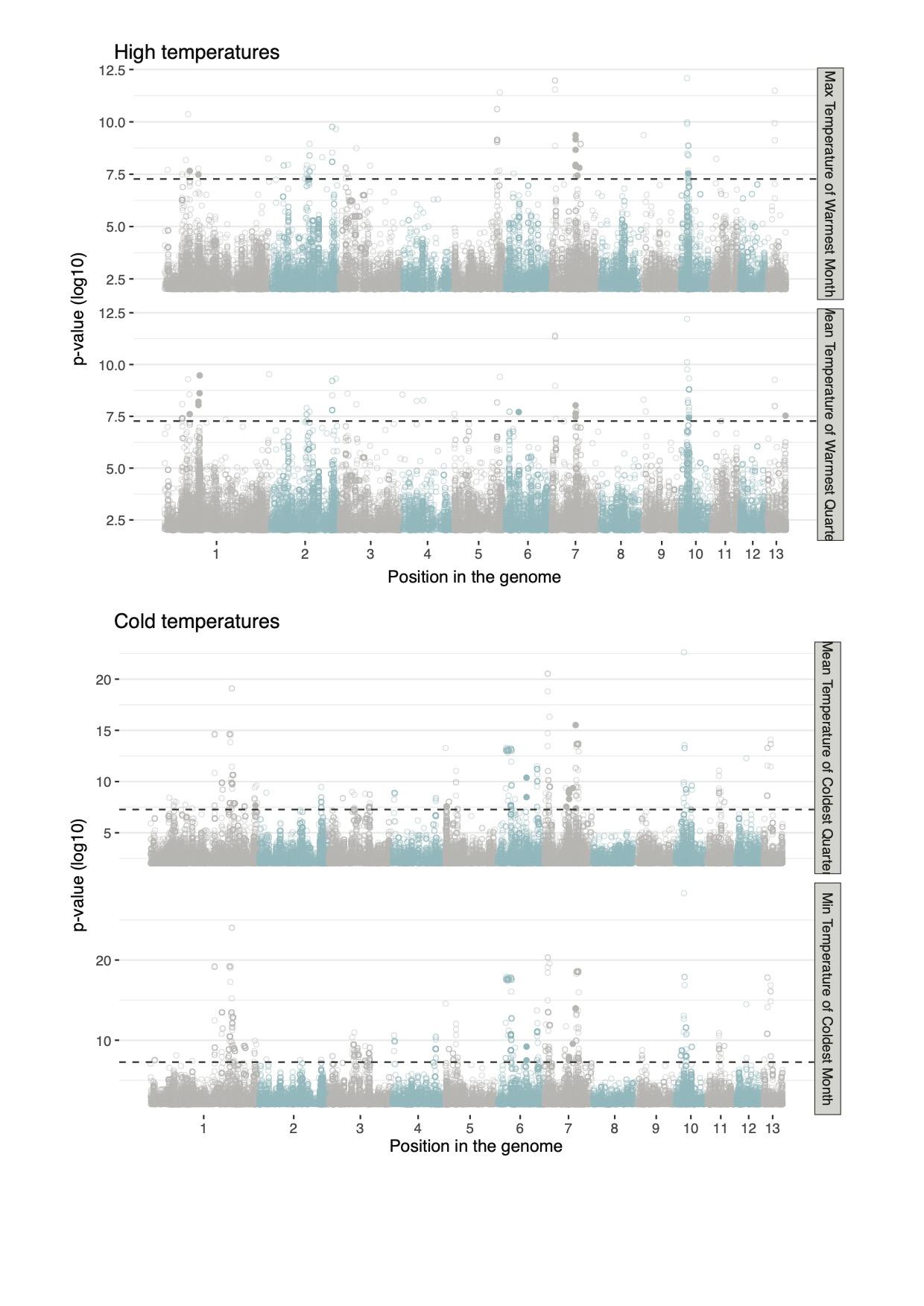
**Figures S14: Manhattan plots of the GEA for high and low temperature variables.**

The x-axis represents the genomic position, each dot represents a short variant with the colors showing alternating chromosomes. The black dashed line is the Bonferonni threshold. Each variant above the threshold is either a filled circle to show a minor allele frequency (MAF) above 5% or an empty circle for lower MAF.

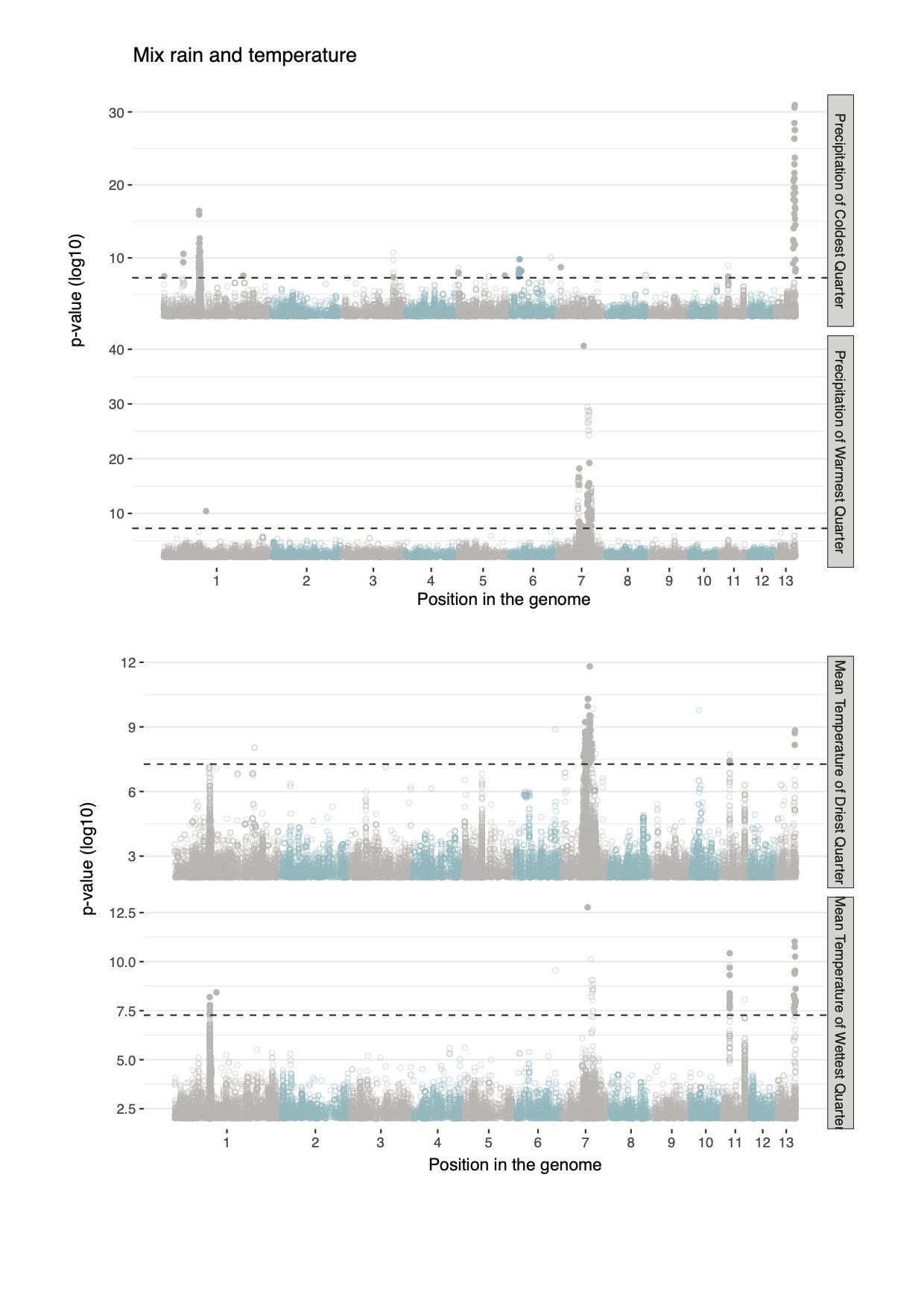

**Figures S15: Manhattan plots of the GEA for bioclimatic variables that include information about both precipitation and temperature.** The x-axis represents the genomic position, each dot represents a short variant with the colors showing alternating chromosomes. The black dashed line is the Bonferonni threshold. Each variant above the threshold is either a filled circle to show a minor allele frequency (MAF) above 5% or an empty circle for lower MAF.

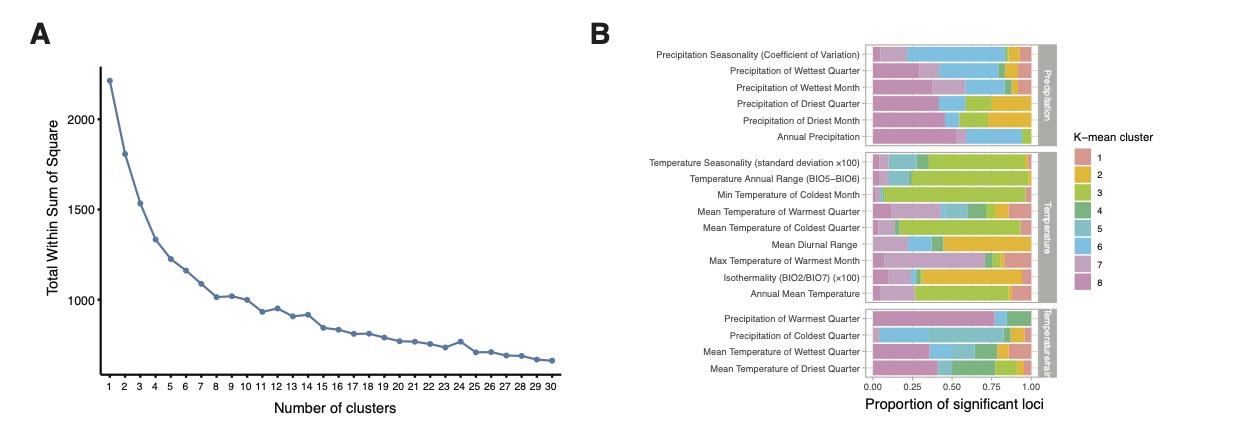

**Figure S16: K-mean clustering of the significant loci based on the presence-absence of the top allele in populations.**

A. Visualization of the total within cluster sum of square, which measures the compactness of the clustering, for a number of clusters ranging between 1 and 30. Using the elbow method, we estimated the best number of clusters at 8.
B. Proportion of variants belonging to either of the 8 clusters (colors) per bioclimatic variable.

**Supplementary tables**

**Table S1**: Isolate metadata, including the isolate name, possible alternative names used in other publications, geographical location of the sampling site and inferred coordinates, sampling year, filtering status (kept or not for the analyses in the manuscript) as well as the bioproject corresponding to the sequencing data.

**Table S2**: Sequencing depth per isolate and per chromosomes, as well as the median on the core chromosomes.

**Table S3**: Clustering results from the population structure. Each line reports the cluster membership value for each cluster and the cluster we assigned it to (based on a 0.75 threshold) or NA if the isolate was considered a hybrid/admixed genotype.

**Table S4**: Bioclimatic variables and their number of significantly associated variants and loci.

**Table S5**: Genes impacted by the significant variants from the GEA analysis.

**Table S6**: Overlap between the GEA significant loci and the QTL from Lendenmann et al.
